# Shannon Entropy as a metric for conditional gene expression in *Neurospora crassa*

**DOI:** 10.1101/2020.10.27.357517

**Authors:** Abigail J. Ameri, Zachary A. Lewis

## Abstract

*Neurospora crassa* has been an important model organism for molecular biology and genetics for over 60 years. *N. crassa* has a complex life cycle, with over 28 distinct cell types and is capable of transcriptional responses to many environmental conditions including nutrient availability, temperature, and light. To quantify variation in *N. crassa* gene expression, we analyzed public expression data from 97 conditions and calculated the Shannon Entropy value for *Neurospora’s* approximately 11,000 genes. Entropy values can be used to estimate the variability in expression for a single gene over a range of conditions and be used to classify individual genes as constitutive or condition-specific. Shannon entropy has previously been used measure the degree of tissue specificity of multicellular plant or animal genes. We use this metric here to measure variable gene expression in a microbe and provide this information as a resource for the *N. crassa* research community. Finally, we demonstrate the utility of this approach by using entropy values to identify genes with constitutive expression across a wide range of conditions and to identify genes that are activated exclusively during sexual development.

## Introduction

Across conditions, individual genes can display expression patterns that can range from conditional to constitutive. When performing Quantitative Reverse Transcription PCR (qRT-PCR) it is crucial to identify constitutively expressed genes for experimental normalization (Huggett *et al.* 2005). Conversely, highly regulated, condition-specific gene promoters are often used in molecular biology to drive conditional expression of a gene under investigation (e.g., an essential gene) or to control expression of reporter genes in certain cell types or environmental conditions (e.g., a gene encoding a fluorescent protein) (Giles *et al.* 1985; Hurley *et al.* 2012; Lamb *et al.* 2013). Moreover, identification of genes that are exclusively expressed during a condition or cell-type of interest can reveal genes that are functionally important. Such genes or promoters are often identified by examining gene expression across just a handful of experimental conditions; however, with the increase in publicly available transcriptomics data it is possible to quantify variation in gene expression across many conditions for a given organism.

In 1963, Claude Shannon laid the basis for information theory, and described the unit known as Shannon entropy (Shannon 1997). A simplistic definition of Shannon entropy is that it describes the amount of information a variable can hold (Vajapeyam 2014). In our case, a variable is a gene, and the information is the collection of expression values from different conditions. If a gene is classified as having low entropy, then the expression values would be generally consistent across different conditions or possess a low amount of information. Instead, if a gene is classified as having high entropy, then the expression of this gene would be highly variable across different conditions and contain a high amount of information.

Since entropy describes information contained in a variable there are a number of uses for such a metric. Previous studies have used entropy to investigate cell and tissue specific expression of genes (Schug *et al.* 2005), identify potential therapeutic targets (Fuhrman *et al.* 2000), characterize periodicity in gene expression (Langmead *et al.* 2002), identify cancerous tissue samples (van Wieringen and van der Vaart 2011), and make genomic comparisons (Machado 2012). Studies using entropy have been carried out in human cell lines (Nathaniel D. Heintzman *et al.* 2009), mouse (Schug *et al.* 2005), plants (Zhang *et al.* 2006), yeast (Timothy R. Lezon *et al.* 2006), bacteria, phage, and metagenomes (Akhter *et al.* 2013) but not yet in filamentous fungi.

*Neurospora crassa* has a 43Mb genome encoding approximately ~11,000 genes (Borkovich *et al.* 2004) (add Nature paper). There is a whole genome knock out collection, and genetic, genomic, and epigenetic studies have been carried out with this organism for more than 100 years (Colot *et al.* 2006). Indeed, *N. crassa* has been used as a model organism for epigenetics, testing fungal enzymes for biomass degradation, and circadian clock studies (Dunlap *et al.* 2007; Tian *et al.* 2009; Aramayo and Selker 2013). As a resource for *N. crassa* researchers, we generated an entropy value for most genes in the *N. crassa* genome using publicly available RNA-seq data, and we validated this approach using previously published lists of housekeeping or inducible genes. This resource has a number of useful applications for the *N. crassa* community.

## Methods

### Public data collection

Entropy calculations were made for all genes in the *N. crassa* genome using public RNA-seq data sets (97 conditions from a total of 173 separate sets including replicates; Table S1).

### Data Analysis

#### Mapping, TPM and entropy calculations

HiSat2 (version 2.1.0) (Kim *et al.* 2019) was used to map all of the SRA accessions to the NC12 genome (NCBI assembly: GCA_000182925.2) using appropriate parameters specific for paired or single end sequence reads (with parameters-RNA-strandness RF or R) to produce bam files which were then sorted and indexed using SAMtools (version 1.3) (Li *et al.* 2009). If experiments contain replicates, the replicate bam files were merged together before obtaining counts with featureCounts from Subread (version 1.6.2) (Liao *et al.* 2014). FeatureCounts was used with parameters-T exon to generate all counts at the gene level. Counts were imported into R where we obtained TPM using the function calculateTPM from the R package scater (McCarthy *et al.* 2017). This package takes in feature-level (in our case, gene-level) counts and gene lengths and outputs the TPM values for each gene. TPM values were then used to calculate the Shannon entropy using the R package BioQC (Zhang *et al.* 2017). The function entropySpecificity was used to calculate the entropy values for all genes in the genome. To examine specific genes sets, we converted from NCU accession numbers to gene identifiers from NCBI Genome Assembly NC12 (GCA_000182925.2) and plotted the kernel density estimation with rug plots.

##### Data Availability Statement

All supplementary tables have been uploaded to Figshare. Table S1 contains SRA accession numbers, short descriptions, total reads, and mapped reads for each public data set used.Calculated entropy values for all *N. crassa* genes are listed in Table S2. Lists of all *N. crassa* genes used to benchmark the entropy values and generate panels in figure 2 and 3 are included in Table S3. Code used to generate the data in this manuscript is available through github. https://github.com/ajcourtney/entropy

## Results and Discussion

Shannon entropy values are useful in measuring the amount of variation in expression levels across different tissues or growth conditions. In order to calculate Shannon entropy values for all *Neurospora crassa* genes, we first compiled a list of available RNA-seq data sets present in the NCBI sequence read archive (SRA) (Table S1). We selected datasets that were generated with the wild type strain Oak Ridge strain background, but we used both mating types. To calculate accurate entropy values, we needed to gather many observations of gene expression across different conditions. We searched the SRA database (Leinonen *et al.* 2011) for *N. crassa* RNA-sequencing entries that were processed at different developmental stages or grown under different conditions. In total we gathered 173 accessions, which represent 97 developmental or growth conditions. We then developed a pipeline to generate entropy values for each gene (Figure 1A). Calculated entropy values are available in Table S2. We first mapped to the NC12 *N. crassa* genome using HiSat2 (Kim *et al.* 2019) to generate bam files. The bam files were then used to generate read counts for each gene in each condition using featureCounts (Liao *et al.* 2014), which assigns reads to genomic features. Once the count file was created, we calculated normalized expression values using the Transcripts per Million (TPM) normalization method to create a matrix of normalized expression values for all genes in all conditions. We then used this expression matrix to calculate the Shannon entropy value for each gene (Zhang *et al.* 2017). This generated entropy values for 10,300 out of 10,398 genes. The remaining 98 genes had 0 read counts in all conditions, so we were unable to calculate entropy. Our final entropy values range from 0.0506 to 6.599. 70% of the genes in the genome possess low entropy values between 0.05 and 1 (7,180/10,300) (Figure 1B). These values include the constitutively expressed genes in the genome. Entropy values above one represent only 30% of the genome (3,120/10,300), corresponding to genes with more condition-specific expression patterns.

**Figure 1:**
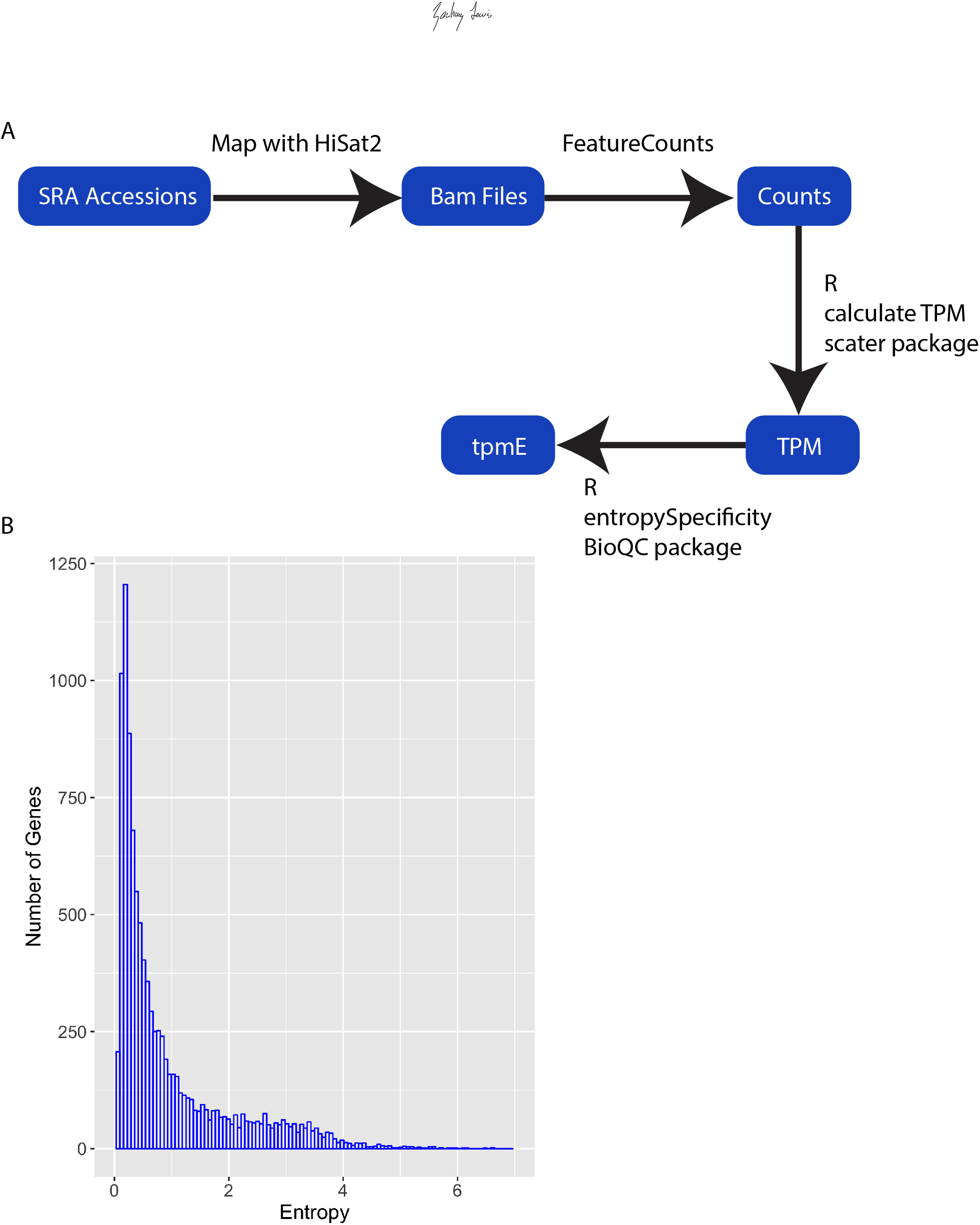
Calculation of Shannon entropy for *N. crassa* genes using public RNA-seq data. A) Schematic of our computational pipeline for calculating Shannon entropy from publically available datasets. B) *N. crassa* genes display a broad range of entropy values. The histogram shows entropy values for all genes. The y-axis is the number of genes found in each bin. The x-axis shows the binned entropy values.

### Validation of entropy as a measure of gene expression variation in *N. crassa*

In order to determine if entropy values are a reliable predictor of expression variability in a microbe, we examined the entropy values generated here for published gene sets expected to be enriched for constitutively expressed genes, or conversely, for gene sets expected to contain genes with highly condition-specific expression patterns. If entropy value is a reliable measure of gene expression variation across conditions, housekeeping genes should be enriched for genes with low entropy values, whereas sets of conditionally-induced genes are expected to be enriched for high entropy values. Two previous studies identified genes useful for RT-qPCR controls in *N. crassa*. One of which published a list of 38 genes classified as “housekeeping genes” based on previously generated microarray and RNA-seq datasets under three different conditions (quinic acid (QA) induction, circadian gene expression profiling, and light response) (Hurley *et al.* 2015), and the other study identified four genes by using previous transcriptomic studies and genes used in related organisms to generate candidates that were validated by quantitative PCR under different conditions (Cusick *et al.* 2014) (Table S1). To visualize the distribution of entropy values in this set of 42 “housekeeping” genes, we plotted a kernel density estimation (KDE) of entropy values (Figure 2A). The KDE is a smoothed version of a histogram estimated from the underlying data. As expected, the highest density of data points in the housekeeping data set is around 0.25 (low entropy) and the density falls sharply around 0.75 (Figure 2A). Two genes in this set possess entropy values above 1.6 and they encode an exo-beta-1,3-glucanase and a UDP-glucose dehydrogenase. We plotted a heatmap depicting TPM values for each gene in each condition with genes ranked by entropy values from low to high (top to bottom) (Figure 2B). Genes with higher entropy values showed significant induction of gene expression under certain conditions, whereas genes with low entropy values displayed consistent expression values across all conditions. In particular, the two genes with high entropy values showed marked induction under certain conditions. Thus, these data highlight the value of performing a comprehensive analysis of conditional gene expression when selecting constitutive control genes.

**Figure 2:**
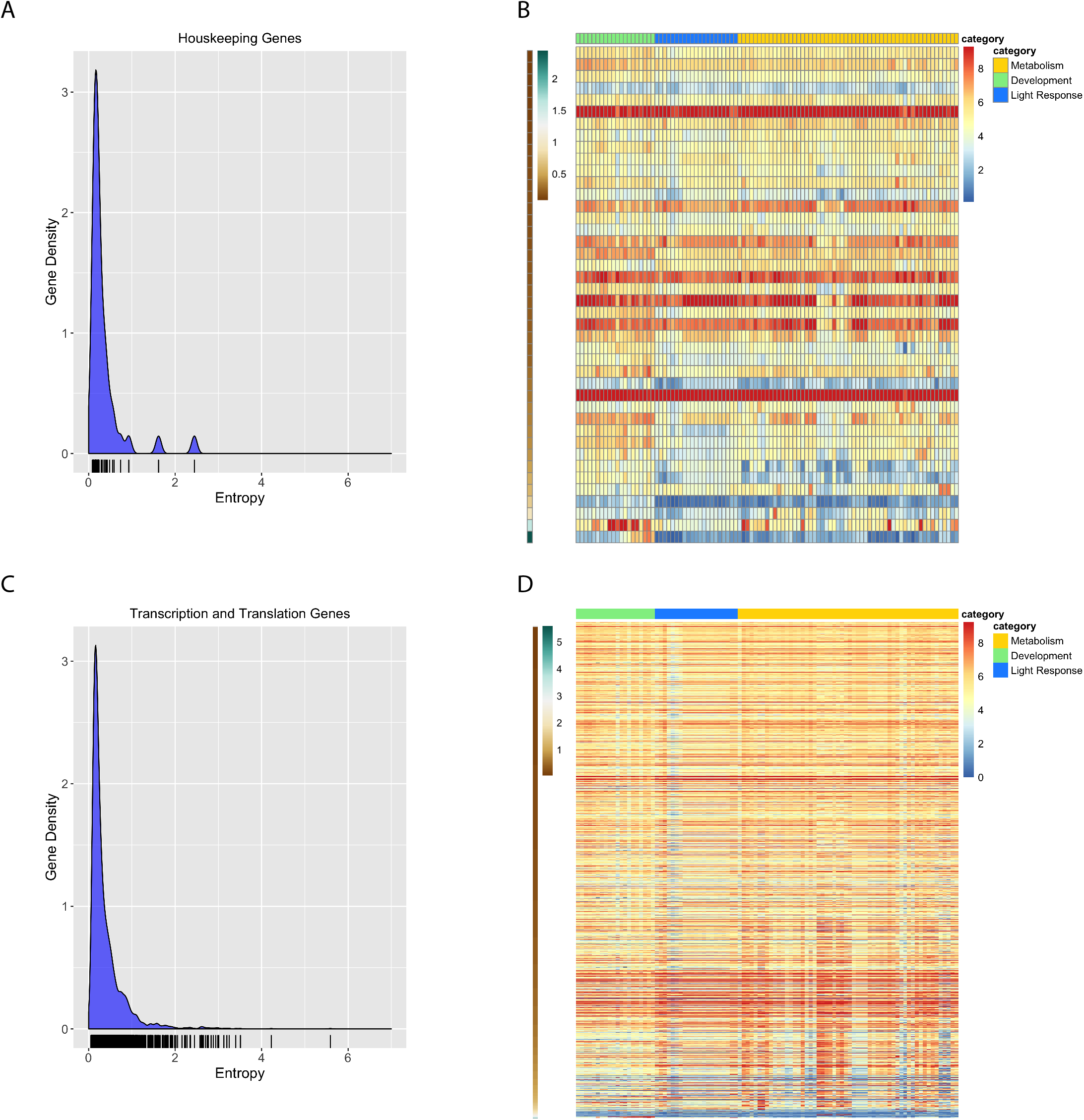
Constitutively expressed genes are characterized by low entropy values. A) The relative frequency of entropy values for a list of housekeeping genes is shown as a kernel density estimation (KDE) plot. The rug plot, black lines on the bottom in the KDE plot represents the individual data points that create the estimation. The y-axis is the probability density, which is the probability for each unit (gene) on the x-axis. The total area below the KDE curve integrates to one. B) The heatmap shows the expression value for housekeeping genes across all conditions analyzed. The expression level for each gene is plotted as the log_2_ transformed transcript per million (TPM) value. Genes (rows) are plotted in ranked order based on the entropy value from low (top) to high (bottom). The scale on the left indicates entropy values for each gene. Each condition (column) has been assigned a category: Metabolism (gold), Development (green), or Light Response (blue). The categories are represented at the top of the heatmap in the three different colors. C) The relative frequency of entropy values for a list of genes related to transcription and translation is shown as a kernel density estimation (KDE) plot. The rug plot, black lines on the bottom in the KDE plot represents the individual data points that create the estimation. The y-axis is the probability density, which is the probability for each unit (gene) on the x-axis. D) Heatmap of log_2_ transformed TPM values from all transcription and translation related genes (rows) ranked by entropy (low to high). Entropy values are depicted by the brown to green heatmap on the left, where brown is low (top) and green is high (bottom). Each condition (column) has been assigned a category: Metabolism (gold), Development (green), or Light Response (blue). The categories are represented at the top of the heatmap in the three different colors.

We further validated the use of entropy as a measure for constitutive gene expression by using the same approach with a published list of genes 2,624 genes involved in transcription and translation (Table S1), reasoning that genes involved in these essential processes would be expressed at similar levels in all 93 conditions we investigated. (Benz *et al.* 2014). The distribution of entropy values for transcription and translation genes resembles the distribution of entropy values for housekeeping genes where the highest density is concentrated at the low end of entropy values (Figure 2C). Many of the genes that possess entropy values above 1.6 are either hypothetical proteins or genes associated with cellular transport or metabolism. We again examined the TPM values for each gene in this set in a heatmap ranked by entropy from low to high and again find mostly steady expression across conditions (Figure 2D).

We next asked if higher entropy values were associated with conditionally expressed genes. The highest entropy values imply that a gene must only be expressed under specific conditions and may only show expression in one or a few of the conditions in the entire RNA-seq dataset. To confirm that higher entropy values were indeed associated with condition-or tissue-specific gene expression, we created KDE plots for 513 genes induced by light (Figure 3A and Table S1) and 3,259 genes that have expression changes during sexual development (Figure 3C and Table S1) (Wu *et al.* 2014) (Wang *et al.* 2014). In both cases, there is a shift in distribution of entropy values toward higher entropy values compared to “housekeeping” or “transcription and translation” genes. We examined TPM values for each gene in each condition using a heatmap ranked by entropy values from low to high (top to bottom) and find that a majority of genes in each gene set show variable expression across conditions, as expected (Figure 3B, D). Genes that have regulation changes during perithecial (sexual) development also show a shift to the right, but with retention of more low entropy genes than in the light induced gene set (Figure 3C). Plotting the TPM values in an entropy ranked heatmap shows that approximately half of these genes are highly expressed across many conditions and half are variably expressed, corresponding to genes with lower entropy values in the density plot (Figure 3D). This implies that half of these genes are not specific to sexual or vegetative cell types even though they show transcriptional changes throughout development (Wang *et al.* 2014).

**Figure 3:**
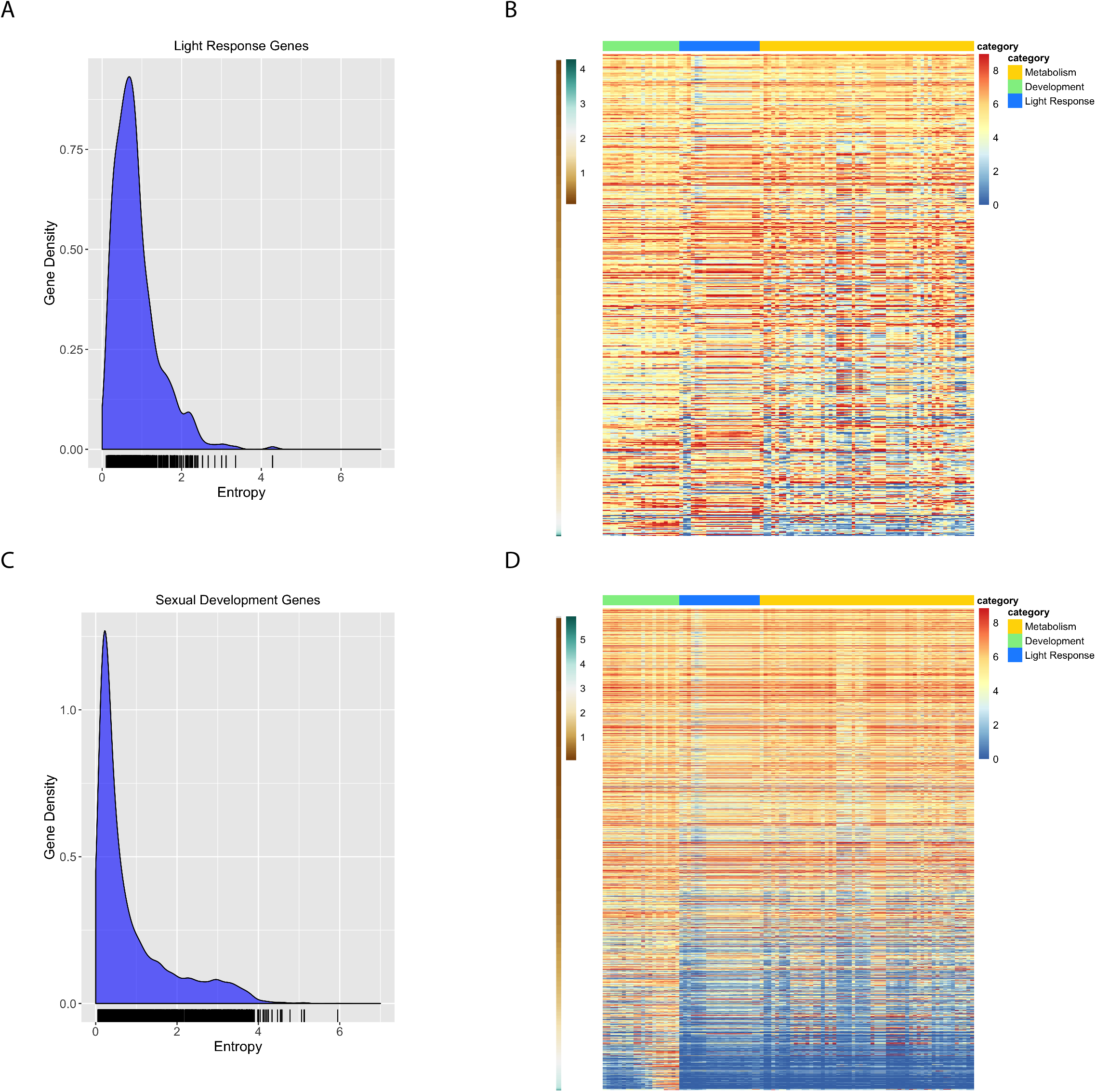
Validating entropy values with previously published light induced genes and genes induced during sexual development. A) The relative frequency of entropy values for a list of light induced genes is shown as a kernel density estimation (KDE) plot. The rug plot, black lines on the bottom in the KDE plot represents the individual data points that create the estimation. The y-axis is the probability density, which is the probability for each unit (gene) on the x-axis. The total area below the KDE curve integrates to one. B) The heatmap shows the expression value for light induced genes across all conditions analyzed. The expression level for each gene is plotted as the log_2_ transformed TPM value. Genes (rows) are plotted in ranked order based on the entropy value from low (top) to high (bottom). The scale on the left indicates entropy values for each gene. Each condition (column) has been assigned a category: Metabolism (gold), Development (green), or Light Response (blue). The categories are represented at the top of the heatmap in the three different colors. C) The relative frequency of entropy values for a list of sexual development genes genes is shown as a kernel density estimation (KDE) plot. The rug plot, black lines on the bottom in the KDE plot represents the individual data points that create the estimation. The y-axis is the probability density, which is the probability for each unit (gene) on the x-axis. The total area below the KDE curve integrates to one. D) The heatmap shows the expression value for sexual development genes across all conditions analyzed. The expression level for each gene is plotted as the log_2_ transformed TPM value. Genes (rows) are plotted in ranked order based on the entropy value from low (top) to high (bottom). The scale on the left indicates entropy values for each gene. Each condition (column) has been assigned a category: Metabolism (gold), Development (green), or Light Response (blue). The categories are represented at the top of the heatmap in the three different colors.

As a final confirmation that entropy can be used as a reliable metric to assess the variation or lack of variation in gene expression levels across many conditions, we plotted the expression levels of 100 genes with the highest entropy values and 100 genes with the lowest entropy values. We took the log_2_ TPM values for all conditions (columns) and plotted them for each gene in a heatmap that was clustered by gene (row) for both the top and bottom 100 genes. As expected, with the lowest entropy values show mostly uniform expression across all conditions (Figure 4A), and genes in the high entropy group displayed highly variable and condition-specific expression (Figure 4B). Together, these data demonstrate that entropy is an effective tool for measuring variation in gene expression levels in a filamentous fungus.

**Figure 4:**
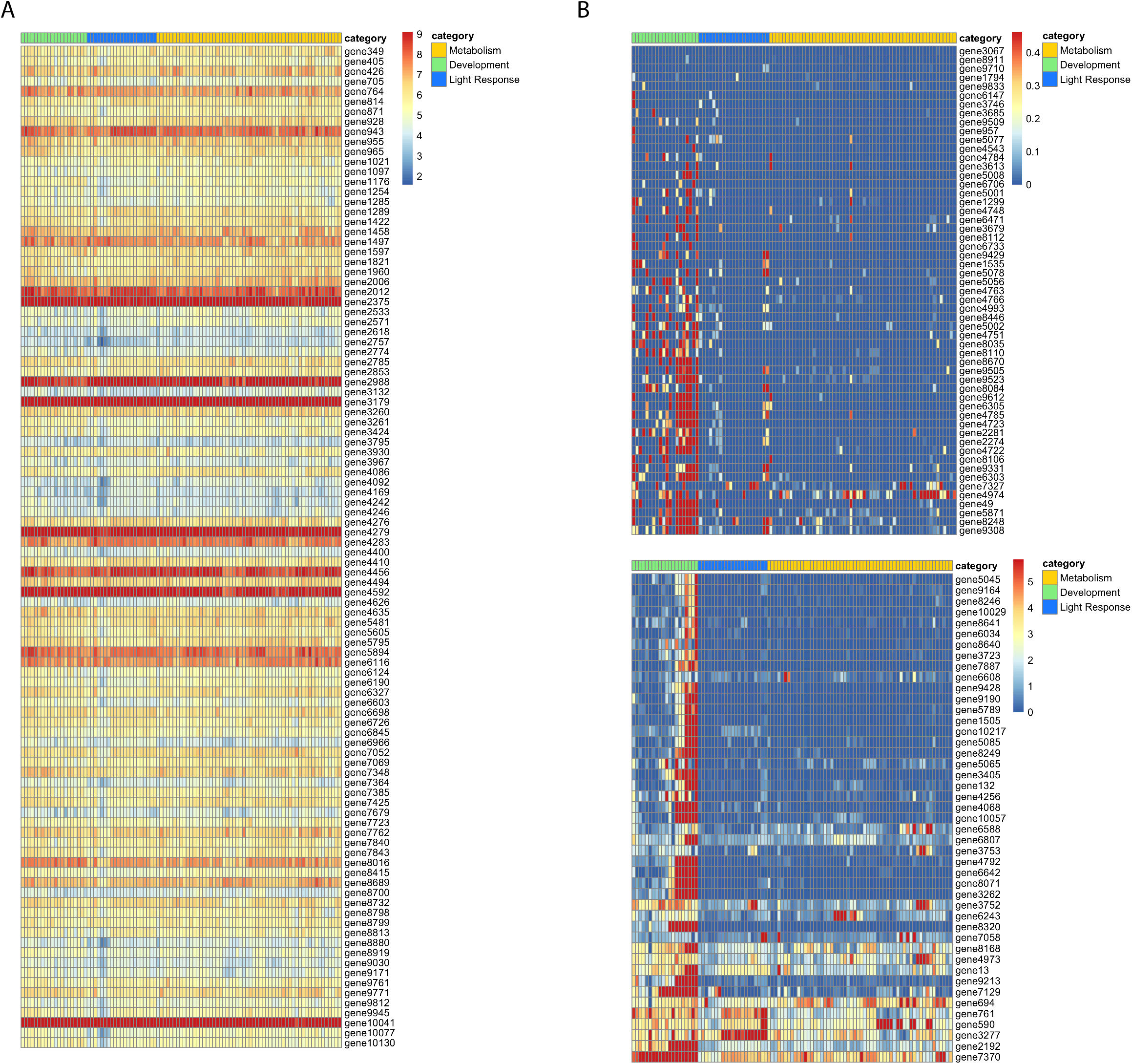
Log_2_ TPM values for highest and lowest ranked genes. A) The heatmap shows the expression values for the 100 genes with the highest entropy values. The expression level for each gene is plotted as the log_2_ transformed TPM value. Each row represents a gene. Gene names are listed on the right side of the heatmap. Each condition (column) has been assigned a category: Metabolism (gold), Development (green), or Light Response (blue). The categories are represented at the top of the heatmap in the three different colors. B) The heatmap shows the expression values for the 100 genes with the lowest entropy values. The expression level for each gene is plotted as the log_2_ transformed TPM value. Each row represents a gene. Gene names are listed on the right side of the heatmap. Each condition (column) has been assigned a category: Metabolism (gold), Development (green), or Light Response (blue). The categories are represented at the top of the heatmap in the three different colors.

The information and code generated in the course of this study could prove useful in a number of ways. First, identifying genes that are induced in a certain condition and display a high entropy value will help identify genes that are condition-specific. In addition, examining entropy values for individual genes can be a useful approach for finding new inducible promoters to use for genetic studies. Condition-specific expressed genes are good starting targets to test for this purpose. The entropy metric determined here can also be used to confirm constitutive expression of genes chosen as controls for RT-PCR. In examining the housekeeping genes from previously published studies it is clear that not all will function as good controls under all conditions, a limitation that was discussed by Hurley and colleagues (Hurley *et al.* 2015). We combined all of their housekeeping genes together, whereas they had them divided into housekeeping genes usable for different conditions in qRT-PCR (QA induction, light response studies, and circadian experiments). Here we can choose genes that will work across all conditions (provided the conditions were represented in the initial dataset). Our approach provides a quantitative metric that can be applied to identify condition-specific genes, as opposed to investigating individual datasets or using controls from previous studies which may not perform as expected. In addition, this methodology is scalable; the initial inclusion of more conditions will only increase the robustness of the metric produced. As more data are published, more datasets can be incorporated. This approach can be used across other fungi in addition to *N. crassa*, provided there are sufficient RNA-seq data publicly available.

## Acknowledgments

We would like to thank Dr. Shannon Quinn for meaningful discussions and insight on this project, and Dr. Kristina Smith for comments on the manuscript. This work was supported by grants from the National Institutes of Health (NIH) (R01GM132644) to Z.A.L. and the National Science Foundation Graduate Research Fellowship Program Grant (DGE-1443117) to A.J.C.

